# The evolution of regulatory elements in the emerging promoter variant strains of HIV-1

**DOI:** 10.1101/2021.04.28.441760

**Authors:** Disha Bhange, Nityanand Prasad, Swati Singh, Harshit Kumar Prajapati, Shesh Prakash Maurya, Bindu Parachalil Gopalan, Sowmya Nadig, Devidas Chaturbhuj, Jayaseelan Boobalan, Thongadi Ramesh Dinesha, Syed Fazil Ahamed, Navneet Singh, Anangi Brahmaiah, Kavita Mehta, Yuvrajsinh Gohil, Pachamuthu Balakrishnan, Bimal Kumar Das, Mary Dias, Raman Gangakhedkar, Sanjay Mehendale, Ramesh Paranjape, Shanmugam Saravanan, Anita Shet, Sunil Suhas Solomon, Madhuri Thakar, Udaykumar Ranga

**Affiliations:** Molecular Biology and Genetics Unit, Jawaharlal Nehru Centre for Advanced Scientific Research (JNCASR), India 560064; Department of Microbiology, HIV Immunology Laboratory, All India Institute of Medical Sciences (AIIMS), India 110029; Division of Microbiology/ Infectious Diseases Unit, St. John’s National Academy of Health Sciences, India 560034; Department of Serology and Immunology, National AIDS Research Institute (NARI), India 411026; Department of Molecular Biology and Genotyping, Y. R. Gaitonde Centre for AIDS Research and Education (YRG CARE), India-600113; Infectious Diseases Laboratory, Y. R. Gaitonde Centre for AIDS Research and Education (YRG CARE), Chennai, India 600113; Department of Clinical Sciences, National AIDS Research Institute (NARI), India 411026; P. G. Hinduja National Hospital and Medical Research Centre, Mumbai, India 400016; Y. R. Gaitonde Center for AIDS Research and Education (YRG CARE), Chennai, Tamil Nadu, India 600113; Department of Medicine, Johns Hopkins University, School of Medicine, Baltimore, Maryland, United States of America

**Keywords:** HIV-1, subtype C, evolution, sequence duplication, latency

## Abstract

In a multicentric, observational, investigator-blinded, and longitudinal clinical study of 764 ART-naïve subjects, we identified nine different promoter-variant strains of HIV-1 subtype C (HIV-1C) emerging in the Indian population, with some of these variants being reported for the first time. Unlike several previous studies, our work here focuses on the evolving viral regulatory elements, not coding sequences. The emerging viral strains contain additional copies of the existing transcription factor binding sites (TFBS), including TCF-1α/LEF-1, RBEIII, AP-1, and NF-κB, created by sequence duplication. The additional TFBS are genetically diverse and may blur the distinction between the modulatory region of the promoter and the viral enhancer. In a follow-up analysis, we found trends, but not significant associations between any specific variant promoter and prognostic markers, probably because the emerging viral strains might not have established mono infections yet. Illumina sequencing of four clinical samples containing a co-infection indicated the domination of one strain over the other and establishing a stable ratio with the second strain at the follow-up time-points. Since a single promoter regulates viral gene expression and constitutes the master regulatory circuit with Tat, the acquisition of additional and variant copies of the TFBS may significantly impact viral latency and latent reservoir characteristics. Further studies are urgently warranted to understand how the diverse TFBS profiles of the viral promoter may modulate the characteristics of the latent reservoir, especially following the initiation of antiretroviral therapy.

**Significance Statement:** A unique conglomeration of TFBS enables the HIV-1 promoter to accomplish two diametrically opposite functions – transcriptional activation and transcriptional silencing. The various phases of viral latency -establishment, maintenance, and reversal -collectively determine the replication fitness of individual viral strains. A profound variation in the TFBS composition of the viral promoter may significantly alter the viral latency properties and the latent reservoir characteristics. Although the duplication of certain TFBS remains a quality unique to HIV-1C, the high-level genetic recombination of HIV-1 may promote the transfer of such molecular properties to the other HIV-1 subtypes. The emergence of several promoter-variant viral strains may make the task of a ‘functional cure’ more challenging in HIV-1C.

## Introduction

Based on phylogenetic association, the viral strains of HIV-1 are classified into four groups (M, N, O, and P), and within group M, into ten different genetic subtypes, A, B, C, D, F, G, H, J, K, L (1), and numerous recombinant forms. Of the various genetic subtypes of HIV-1 unevenly distributed globally, HIV-1C and its recombinant forms are responsible for nearly half of the global infections (2–4). Despite the high prevalence of HIV-1C, only a limited number of studies are available examining the causes underlying the expansion of these viral strains and their impact on disease manifestation.

Although the basic architecture of HIV-1 LTR is broadly conserved among the diverse HIV-1 genetic families, subtype-associated differences are manifested (5). The configuration of transcription factor binding sites (TFBS), including those of NF-κB, NF-AT, AP-1, and other regulatory elements such as the TATA box, and the TAR region, in HIV-1C LTR (C-LTR) differs from that of the other viral subtypes (5). Of the TFBS variations, differences in the copy number and sequence of the NF-κB motif are unique to HIV-1C. HIV-1C LTR typically contains three or four NF-κB motifs in the enhancer region compared to only one motif present in HIV-1A/E or two motifs in all the other HIV-1 subtypes (6). Further, the additional copies of the NF-κB motif in C-LTR are genetically variable, alluding to the possibility of the viral promoter being receptive to a diverse and broader range of cellular signals. For instance, the four copies of the NF-κB motif in the enhancer of 4-κB viral strains represent three genetically distinct NF-κB binding sites (6). Apart from the NF-κB motif, other regulatory elements, including AP-1, RBEIII, and TCF-1α/LEF-1, also show subtype-associated variations, although the impact of such variations has not been examined.

Several publications reported the insertion or deletion of TFBS in HIV LTR. One example is the sequence duplication of the TCF-1α/ LEF-1, RBF-2, AP-1, and c-EBPα binding motif in the modulatory region of the LTR, technically called the most frequent naturally occurring length polymorphism (MFNLP) (7–10). Approximately 38% of the HIV-1B viral isolates contain MFNLP, a phenomenon believed to be a compensatory mechanism to ensure the presence of at least one functional RBEIII site in the LTR (11). Although RBEIII duplication has been found in several subtypes, the significance of this phenomenon has been examined predominantly in HIV-1B. It, however, remains inconclusive whether the presence of RBEIII duplication is directly associated with reduced viral replication or slower disease progression (12). A small number of reports examined RBEIII duplication in HIV-1C infection, however, without evaluating its effect on the replication fitness of the viral strains and disease progression (13).

Over the past several years, our laboratory has documented the emergence of LTR-variant strains of HIV-1C in India and elsewhere (14–16). While the appearance of genetic diversity and such diversity impacting viral evolution are common to the various genetic subtypes of HIV-1, the genetic variation we describe in HIV-1C is non-sporadic and radically different in an important aspect. Viral evolution in HIV-1C appears to be directional towards modulating transcriptional strength of the promoter by creating additional copies of the existing TFBS, such as NF-κB, AP1, RBEIII, and TCF-1α/ LEF-1 motifs, by sequence duplication and co-duplication. A single viral promoter in HIV-1 regulates two diametrically opposite functions critical for viral survival -transcriptional activation and silence. Hence, any variation in the constitution of the TFBS (copy number difference and/or genetic variation), may have a profound impact on viral replication fitness.

Here, in a multicentric, observational, non-interventional, investigator-blinded, and longitudinal clinical study, we examined the promoter sequences of 455 primary viral isolates derived from ART-naïve subjects. We show that the magnitude of TFBS variation is much larger than we reported previously. At least nine different TFBS variant viral strains have emerged in recent years. Using the Illumina MiSeq platform, we attempted to characterize the proviral DNA of a selected subset of viral variants containing the RBEIII motif duplication. The data allude to the possibility that some of the emerging strains could achieve greater replication fitness levels and may establish expanding epidemics in the future, which requires monitoring. This work provides important insights into the HIV-1 evolution taking place at the level of population in India.

## Results

### The magnitude of TFBS variation in HIV-1C LTR

We collected 764 primary clinical samples from four different clinical sites in India, All India Institute of Medical Sciences (AIIMS), New Delhi; National AIDS Research Institute (NARI), Pune; St. John’s Medical Hospital, Bangalore; and Y. R. Gaitonde Centre for AIDS Research and Education (YRG CARE), Chennai, between 2017-2019. Using the genomic DNA, we determined the sequence of the U3 region in the LTR of 520 of 764 viral samples, whereas the amplification of the rest of the samples failed. The genetic typing of 455 of 520 sequences could be accomplished successfully. The proportion of viral variants across the four clinical sites was comparable without geographic skewing (*SI Appendix*, Table S3). The large majority of the variant LTRs contain additional copies of the existing TFBS, including that of NF-κB, RBF-2, AP-1, and TCF-1α/LEF-1 binding sites (Fig. 1, *SI Appendix*, Fig. S1). Based on the TFBS profile, the sequences of the copied TFBS, and their temporal location, the various viral promoters may be classified into three categories.

**Fig. 1.**
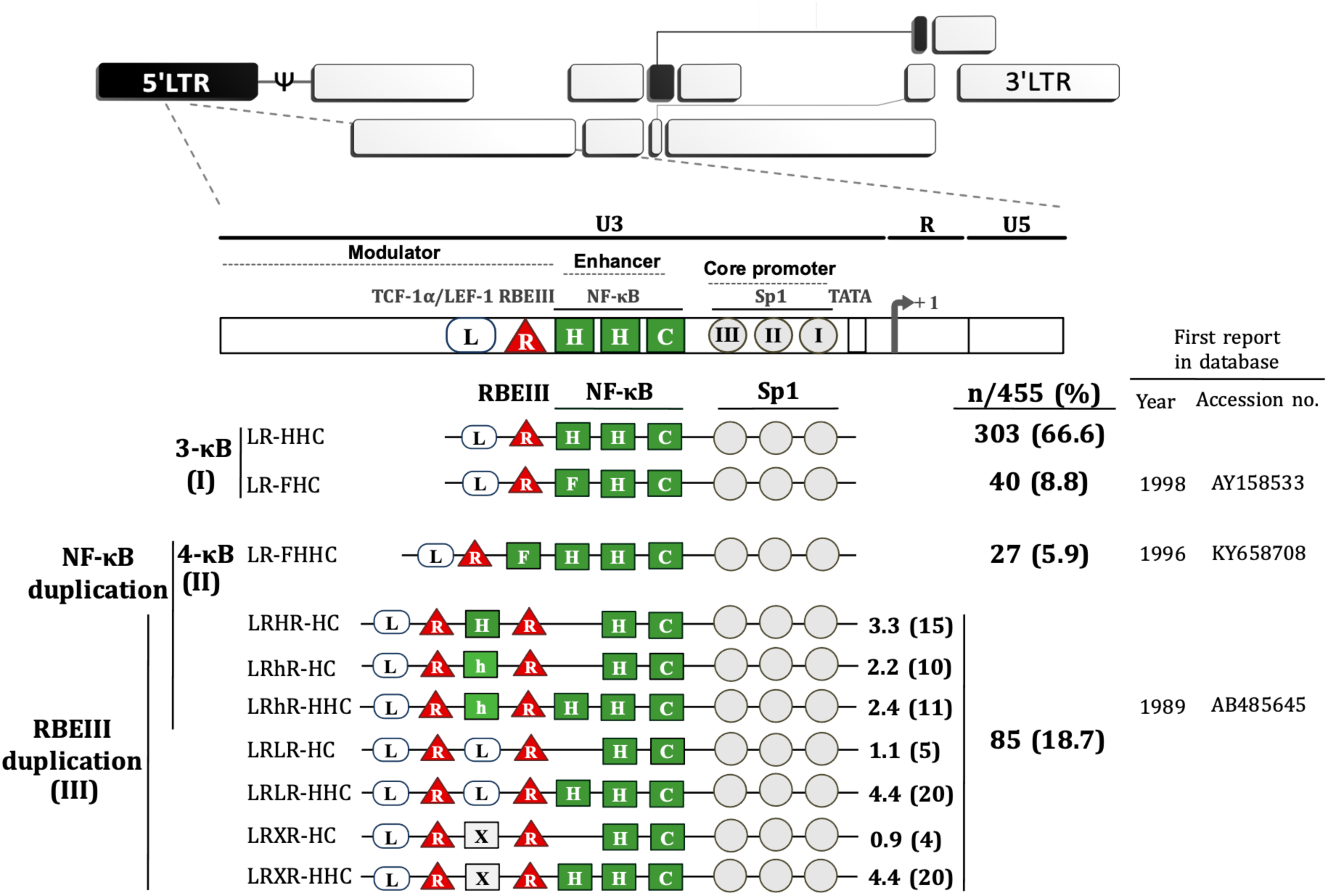
The magnitude of TFBS variation in HIV-1C LTR. The upper panel represents genome organization of HIV-1 followed by the TFBS arrangement in the canonical HIV-1C LTR. Sp1 motifs are depicted as grey circles, RBEIII motifs (R) as red triangles, and TCF-1α/LEF-1 sites (L) as open rectangular boxes. The various types of NF-κB binding sites (H, C, F, and h) are depicted as green square boxes. The various HIV-1C viral strains are classified into three main categories based on the NF-κB and/or RBEIII motif duplication. (I) The 3-κB LTR viral strains. The canonical viral strains (LR-HHC) continue to remain the major category. The LR-FHC strains contain three different NF-κB motifs, and these strains represent a new variant identified for the first time. (II) The canonical 4-κB LTR viral strains. The LR-FHHC-LTR contains four tandem copies of NF-κB motif in the enhancer and its frequency appears to be dropping since the previous reports (Bachu M. et. Al., 2014). (III) The viral strains containing the RBEIII site duplication. The two RBEIII sites are separated by an interceding sequence that constitutes an additional copy of a κB-motif (H), κB-like motif (h), TCF-1α/LEF-1 motif (L) or sequence without a distinct pattern (X). The analysis is from 455 samples as we could not type 65 of 520 LTR sequences.

Category-1 is represented by the viral promoters containing three NF-κB motifs without any other TFBS duplication. This group consists of the canonical LR-HHC-LTR and a new variant LR-FHC-LTR (see below). The canonical LR-HHC-LTR represents the largest group among all HIV-1C promoters, comprising 303 of 455 sequences (66.6%). This LTR contains three tandemly arranged NF-κB sites in the enhancer representing two distinct κB-motifs, two H-κB motifs (5’-GGGACTTTCC-3’), and one C-κB motif (5’-GGG<u>G</u>C<u>G</u>TTTCC-3’, differences underlined). Immediately upstream of the viral enhancer, an RBF-2 binding site (R, RBEIII motif, 5’-ACTGCTGA-3’) and further upstream a TCF-1α/LEF-1 site (L, 5’-TACAAA/GG/A-3’) is located. The canonical HIV-1C promoter is identified here as LR-HHC-LTR to denote the specific arrangement of the three categories of TFBS from 5’ to 3’. The second member of the group contains a variant LR-FHC-LTR comprising 40 of 455 (8.8%) sequences. The characteristic feature of these viral strains is the presence of three genetically distinct NF-κB motifs in the viral enhancer. Thus, the two viral strains of category-1 formed the major proportion of all the LTRs, 343/455 (75.4%) (Fig. 1). Multisequence alignments of several representative viral strains of the canonical LR-HHC (Fig. 2*A*) and the variant LR-FHC (Fig. 2*B*) are presented. The LR-FHC-LTR, being reported here for the first time, may have originated from the 4-κB viral strain FHHC of category 2 described below.

**Fig. 2.**
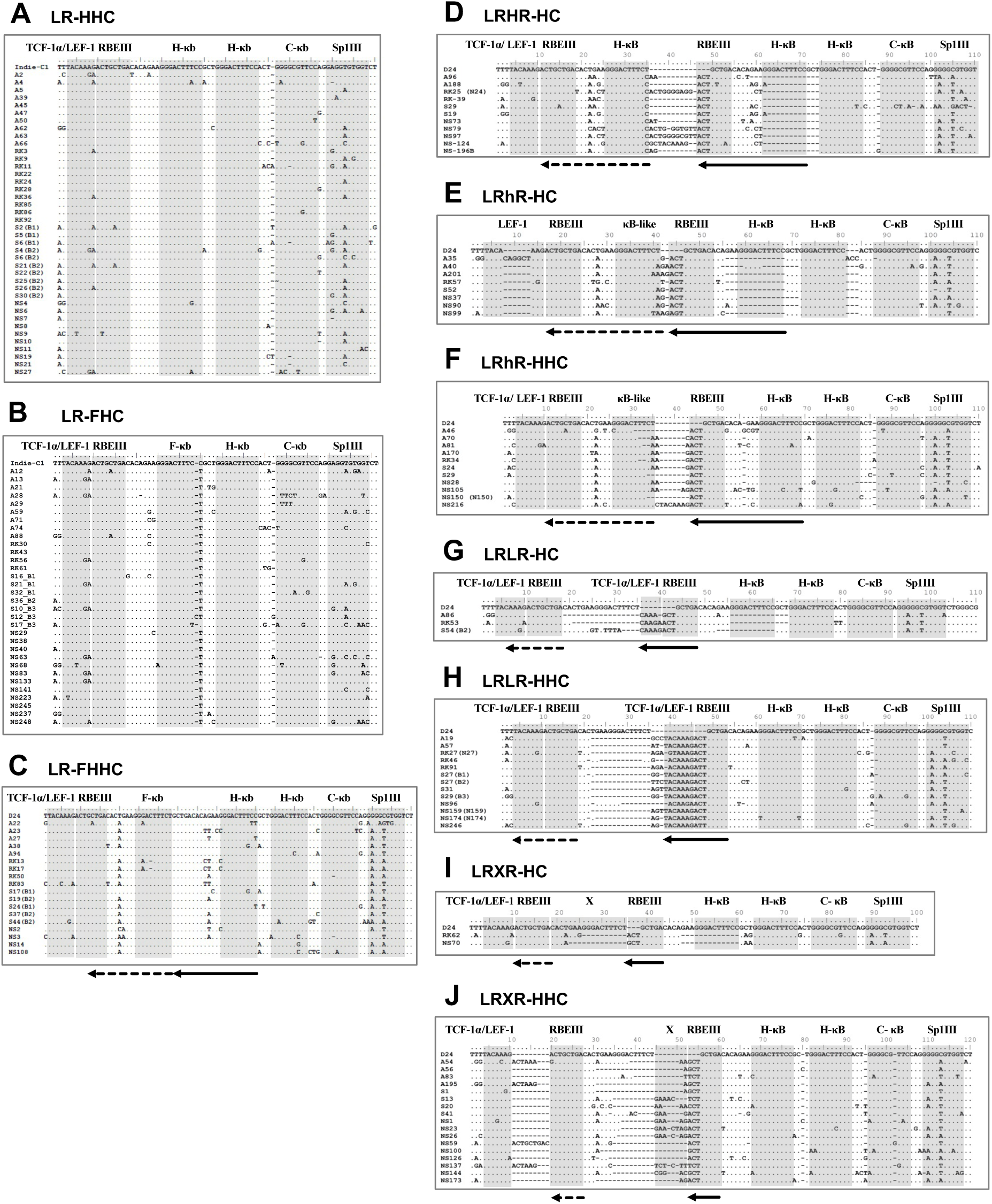
Multiple sequence alignment of the LTR sequences of HIV-1C strains. The alignment contains a few representative sequences under each category. Patient identity is depicted on the left side of alignment. TFBS of significance are highlighted using grey shade boxes and labelled on the top. (A) The LR-HHC (the canonical 3-κB strains) sequences are aligned with the Indie C1 (AB023804.1) reference sequence. (B) The alignment of the LR-FHC variant viral sequences (C) The alignment of the LR-FHHC (4-κB variants) sequences with the D24 (EF469243.2 reference sequence. (D, E, F, G, H, I and J) Sequence alignment of seven types of double-RBEIII variant viral strains. The intervening sequences between the two RBEIII sites represent an H-κB (H), κB-like (h), hLEF (L), or un-typable (X) motif. The solid and dotted arrows represent original and duplicated sequence, respectively.

Category-2 viral LTRs contain an additional (fourth) NF-κB binding site (F-κB site, 5’-GGGACTTTC<u>T</u>-3’) located downstream of the RBEIII site. The 4-κB LTRs thus, contain one F-, two H-, and one C-κB motifs, in that order, hence are labeled LR-FHHC. An alignment of several representative 4-κB viral sequences shows a high degree of sequence conservation at all the TFBS (Fig. 2*C*).

### Double RBEIII LTRs show profound variation in number, genetic sequence, and position of TFBS

Category-3 viral strains, representing 18.7% (85/455) sequences analyzed here, are characterized by the duplication of the RBEIII site in the modulatory region upstream of the viral enhancer, which is analogous of MFNLP described previously in HIV-1B (Fig. 1). Several unique molecular properties qualify the duplication of the RBEIII site as described below.

Firstly, the duplicated sequence consists of three different elements -two of the elements invariably co-duplicated, with the third element being variable. The two elements co-duplicated are the eight base RBEIII core motif (5’-ACTGCTGA-3’) which binds the RBF-2 factor, and a down-stream seven base motif (5’-TGACACA-3’) that forms a binding site for a heterodimer of c-Jun and ATF (*SI Appendix*, Fig. S1). Notably, the 3’-TGA residues of the RBEIII core sequence overlap with the binding site of the c-Jun: ATF heterodimer. Thus, a part of the duplicated sequence (5’-**ACTGC<u>TGA</u>**<u>C</u>*ACA*-3’, the bolded sequence binds RBF-2, the underlined sequence is expected to bind c-Jun, and the italicized sequence ATF) appears to mediate the binding of a complex of transcription factors consisting of RBF-2, c-Jun, and ATF. Notably, the entire sequence consisting of the eight base core RBF-2 binding motif, overlapping c-Jun, and ATF binding motif is strictly conserved in the RBEIII motif duplication. Secondly, in addition to the duplication of the 5’-ACTGCTGACACA-3’ sequence, additional sequences are also co-duplicated, forming the basis for further classification of the viral strains. In a canonical HIV-1C LTR, the RBEIII motif is flanked by an upstream TCF-1α/LEF-1 site (5’-TACAAA/GG/A-3’) and a downstream NF-κB element (5’-GGGACTTTCC-3’). When the RBEIII motif is duplicated, one of these two TFBS is also co-duplicated, forming at least two sub-groups -viral strains containing a TCF-1α/LEF-1 or NF-κB site co-duplication. Thirdly, in the variant LTRs, the two RBEIII sites are not contiguous but are invariably separated by an intervening sequence that usually forms a binding site for NF-κB or TCF-1α/LEF-1, as described above.

Based on the nature of the intervening sequence, the viral strains may be classified into two major subgroups, one containing the presence of a canonical NF-κB (5’-GGGACTTTCC-3’, LRHR-HC), or κB-like motif (5’-GGGACTTTC<u>A</u>-3’, which is denoted as ‘h’ κB-motif in the present work, strains LRhR-HC and LRhR-HHC) and the other a TCF-1α/LEF-1 (5’-TACAAA/GG/A-3’, LRLR-HC and LRLR-HHC) or an incomplete TCF-1α/LEF-1 binding sequence. A third sub-group may also be identified where the intervening sequence does not seem to form a binding site for a defined host factor (LRXR-HC and LRXR-HHC). Multisequence alignments of the variant viral promoters are presented (Fig. 2*D-J*). Of note, the three NF-κB motifs in LRHR-HC, LRhR-HC, and LRhR-HHC strains are not arranged in tandem, thus, blurring the distinction between the viral enhancer consisting of the only NF-κB motifs and the upstream modulatory region.

In a phylogenetic analysis of 461 LTR sequences, compared with 33 reference sequences, all the viral strains of the cohort grouped with HIV-1C ascertaining their genetic identity (Fig. 3). A single sequence A105-9811 is the only exception that clustered with HIV-1K reference sequences. Notably, the various viral sequences combined homogeneously without forming separate clusters based on the TFBS variation.

**Fig. 3.**
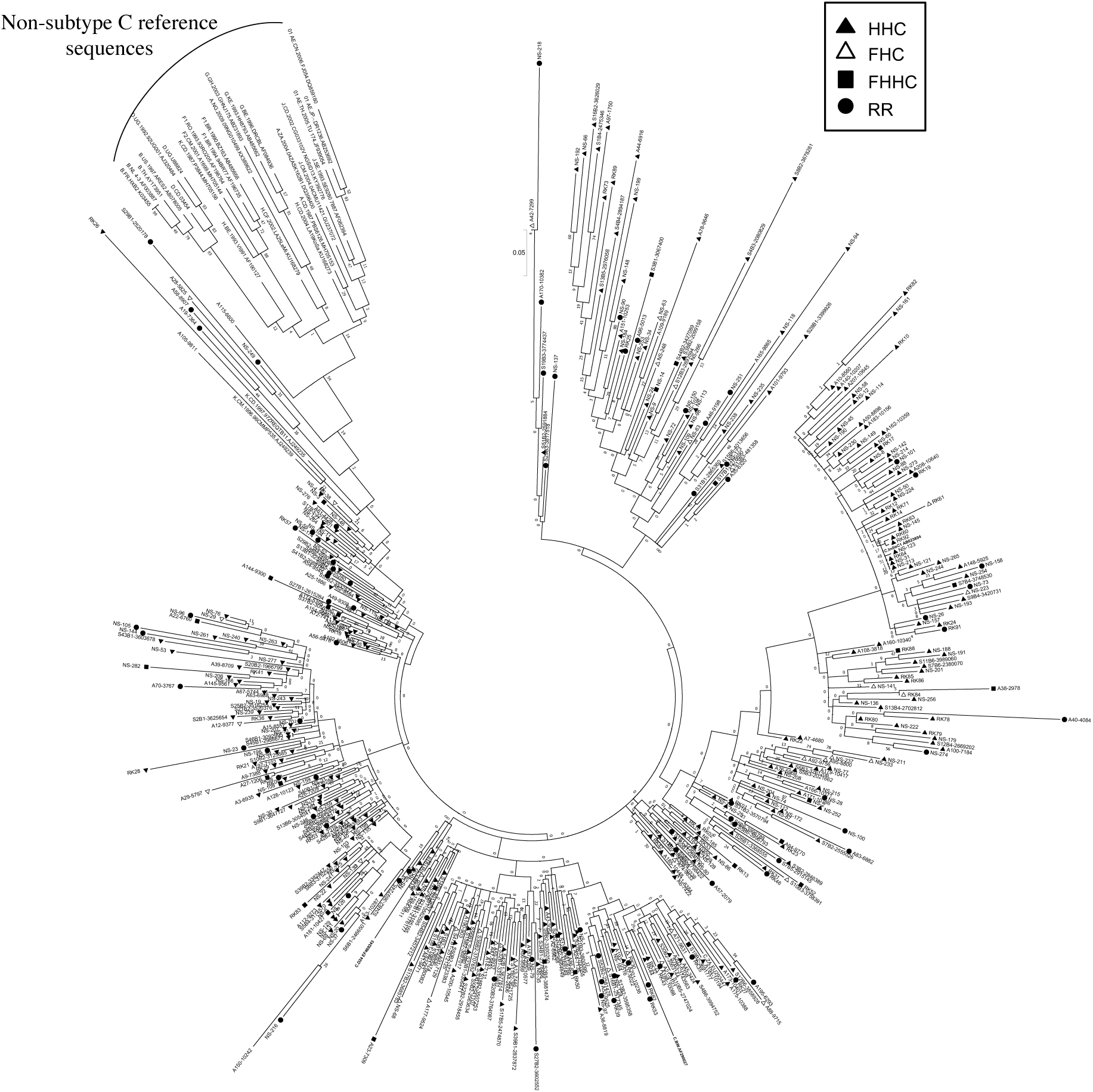
Phylogenetic analysis of HIV-1C LTR variant sequences. A total of 461 viral sequences isolated from study participants are included in the analysis. The analysis also includes four HIV-1C reference sequences and three sequences representing each major genetic subtype of HIV-1 as described in materials and methods. All the reference sequences are represented in bold. Different LTR variant types are represented using different symbols as depicted. The analysis was performed with 1000 bootstrap and the percentage of clustering are shown. The tree is drawn to scale with branch lengths in the same units as those of the evolutionary distances used to infer the phylogenetic tree and the scale is shown at the beginning of the tree. There were a total of 412 positions in the final dataset. Evolutionary analyses were performed using MEGA7.0 software.

### Longitudinal analysis of prognostic markers

Increased transcriptional strength of the LTR may augment plasma viral load modulating various prognostic markers and immune activation markers. To this end, we monitored a few prognostic markers, such as the plasma viral load, CD4 cell count, and soluble CD14 levels, in the blood samples at the baseline and at two or three follow-up time-points spaced six months apart. Unfortunately, this objective could be fulfilled only partially given practical constraints. Following the primary screening and typing of the viral promoters, we could recruit only 208 of the 455 study participants for the follow-up analysis. Subsequently, with the implementation of the ‘Test and Treat’ policy in 2017 in India, several participants enrolled for the follow-up were excluded from the study. Consequently, the longitudinal analysis could be accomplished only with a small number of study participants, who did not prefer to switch to ART.

The demographic features of the 208 study participants at the baseline are summarised (Table 1). Of the 208 study participants, the percentage of female, male, and transgender are 55.3% (115/208), 43.8% (91/208), and 1.0% (2/208), respectively. The average (mean) age of the study participants was 34.4±8.45 years (median = 33 years). For the subsequent analyses, all the viral strains were classified into four categories based on the nature of the TFBS variations identified in the viral promoter – HHC (the conventional LTRs containing the HHC κB-binding sites), FHC (all the three κB-binding sites are genetically distinct in these LTRs), FHHC (the LTRs contain four κB-binding sites), and RR (double-RBEIII strains; the seven viral variant strains are pooled into a single category, given the limited number of samples in individual groups).

**Table 1.** Characteristics of study participants in the longitudinal study at the time of recruitment

| Parameters |  |
| --- | --- |
| Total number of participants | 208 |
| Gender |  |
| Female | 55.3% (n=115) |
| Male | 43.8% (n=91) |
| Transgender | 1.0% (n=2) |
| Age |  |
| Mean and SD | 34.4±8.45 |
| Median | 33 |
| ART status | Naïve |

We compared the levels of plasma viral load, CD4 cell count, and sCD14 among these four categories at the baseline in a cross-sectional analysis. This analysis did not show a significant difference in any of the three parameters among the four groups (Fig. 4, left panels). The median values of all the parameters were comparable among the four groups. At M0, the median PVL values for the HHC, FHC, FHHC and, RR groups were 12,609.0, 13,553.0, 10,440.0, and 6,321.0 copies/ml, respectively (Fig. 4, left panel). The median values of CD4 cell count and sCD14 were also similarly comparable among the four groups. We also compared the three clinical parameters between the baseline and 12 M time-point for PVL and sCD14 and baseline, 6 M and 12 M for CD4 cell count (Fig. 4, right panels). All the three parameters appeared to remain stable without a significant change between the baseline and 12 M. The promoter configuration did not appear to make a significant difference for any of the three parameters examined. For instance, the median PVL values at 12 M were 18,450.0, 52,539.0, 13,266.0, and 6,090.0 copies/ml, for HHC, FHC, FHHC, and RR groups, respectively. The CD4 cell count remained stable over the 12-month observation period among all four groups. The sCD14 levels appeared to show an increasing trend among all four groups at the follow-up; these differences, however, were not statistically significant. Of note, in complete-case analysis, a trend of low-level plasma viral load was manifested for the RR variants compared to the three other groups. However, the viral load increased between the time points among all the groups (*SI Appendix*, Fig. S3*A*).

**Fig. 4.**
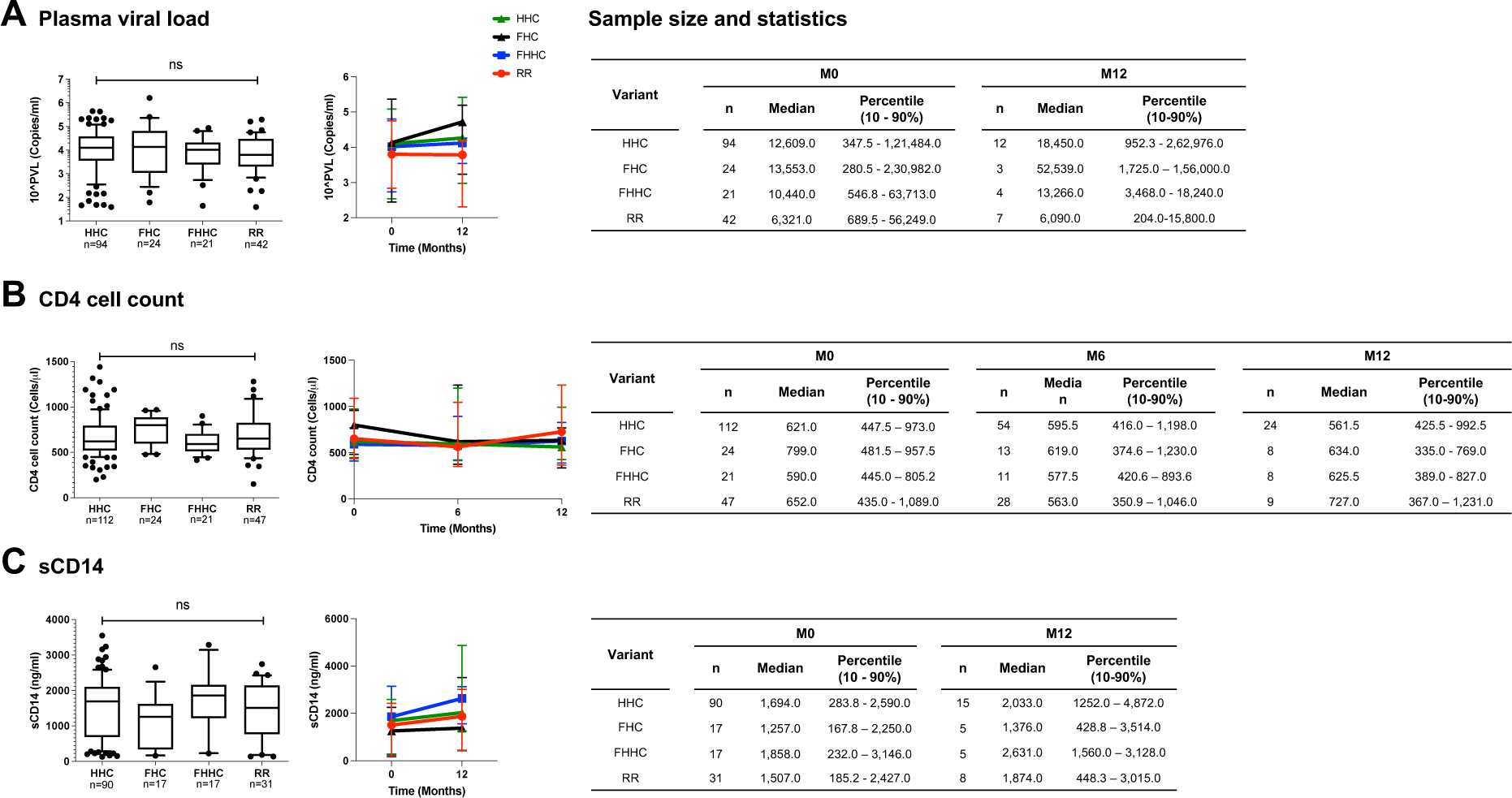
Cross-sectional and longitudinal analysis of prognostic markers. Plasma viral load (A), CD4 cell count (B), and soluble CD14 (C) levels among the four study arms are presented at the baseline (left panels) and follow up points (right panels). The number of samples included under each evaluation are presented in the tables. Given the limited sample numbers, several groups were pooled under the double-RBEIII arm. A non-parametric test i.e. Kruskal-Wallis test was applied for the statistical analysis of the plasma viral load. One-way ANOVA analysis was applied to CD4 count and sCD14.

### The coexistence of viral variants in natural infection

Of the various promoter variant viral strains described in the present work, the emergence of LTRs containing RBEIII duplication is relevant to HIV-1 latent reservoirs. In the context of HIV-1B, the RBEIII site, as well as the AP-1 motif, are known to play a predominantly suppressive role, especially in the absence of cellular activation (17–19). The relative proportion of reads representing single Vs. double RBEIII motif-containing viral sequences in a co-infection may offer leads as per the biological significance of RBEIII duplication in natural infection.

To this end, we identified a subset of four of 85 subjects of our cohort who showed the presence of a co-infection of single- and double-RBEIII viral strains in Sanger sequencing. The clinical profile of the four subjects (2079, 3767, 4084, and VFSJ020) is summarised (Table 2). We performed an NGS analysis, using the MiSeq Illumina platform (*SI Appendix*, Fig. S4), of the whole blood genomic DNA and plasma viral RNA of the four subjects at the baseline and two or three follow up time points.

**Table 2.** The clinical profile of the four study subjects containing a duplication of the RBEIII motif

| Subject ID | Age/<br>Gender | Enrollment<br>date | Promoter<br>variant | PVL (number of RNA copies/ml) | | CD4 (cells/μl) | | | | sCDS14 | |
| --- | --- | --- | --- | --- | --- | --- | --- | --- | --- | --- | --- |
|  |  |  |  | M0 | M12 | M0 | M6 | M12 | M24 | M0 | M12 |
| 2079 | 32/F | 01/06/2016 | LR-HHC and<br>LRLR-HHC | 185 | 9,989 | 989 | 332 | 558 | 1,439 | 1,146.00 | 729.56 |
| 3767 | 40/F | 29/06/2016 | LR-HHC and<br>LRhR-HHC | 3,742 | - | 566 | 292 | 661 | 430 | 139.00 | - |
| 4084 | 38/F | 07/05/2016 | LR-HHC, LR-HC, and LRhR-HC | 6,924 | 5,179 | 509 | 415 | 442 | 613 | 983.00 | 448.28 |
| VFSJ020 | 38/F | 22/09/2016 | LR-HHC and<br>LRhR-HC | 21,200 | - | 461 | - | 579 | 516 | 2,204.43 | - |

The NGS data confirmed the coexistence of single- and double-RBEIII viral strains in all four subjects in both the DNA and RNA compartments. Importantly, in two subjects (2079 and 4084), the single-RBEIII strains represented a significantly larger proportion of reads in the plasma RNA compared to the double-RBEIII strains (Fig. 5). Only in subject VFSJ020, the double-RBEIII reads dominated the single-RBEIII reads at all the time points, whereas in subject 3767 a mixed profile was observed. The data between the replicate samples are consistent with each other ascertaining the reproducibility of the analysis. A broad level concurrence between the plasma RNA and genomic DNA was also noted.

**Fig. 5.**
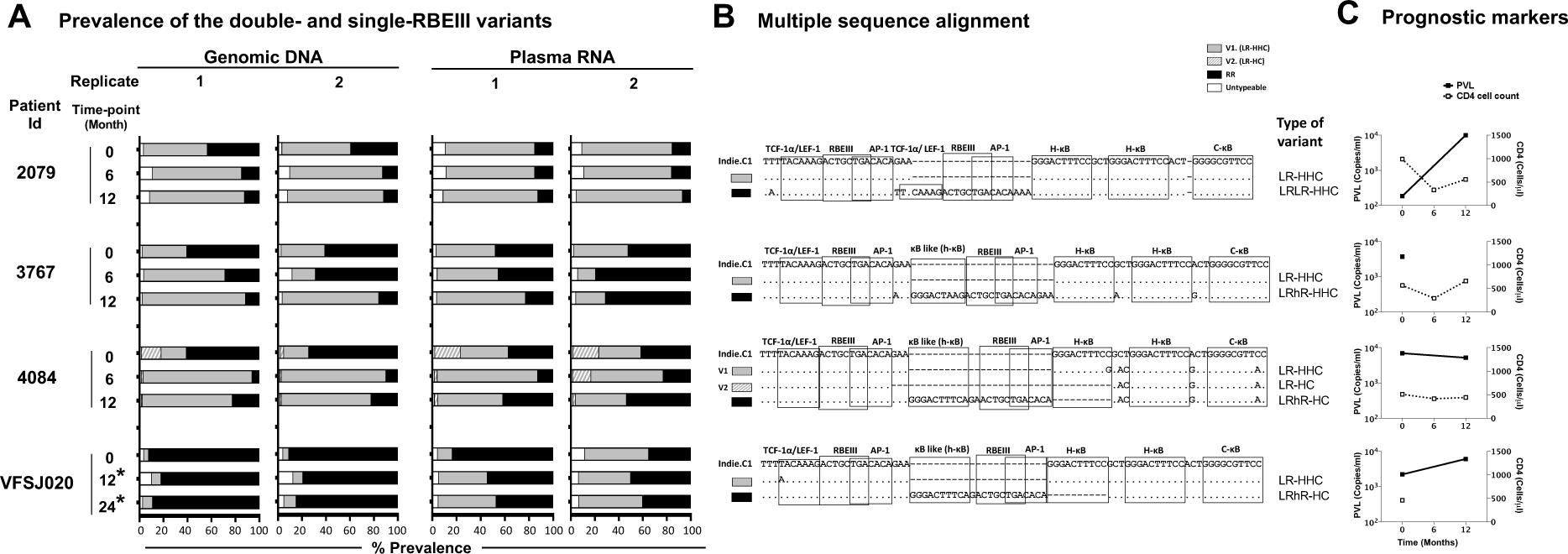
The frequencies of single- and double-RBEIII variants in a subset of study participants. (A) Two independent rounds of analyses were performed (replicates 1 and 2) using both the whole blood genomic DNA and plasma viral RNA. The samples were collected at six-month intervals as shown. An asterisk (*) represents the samples collected post-ART. The dark, grey and hollow bars represent the percentage prevalence of double-RBEIII, single-RBEIII and minority/un-typable viral strains, respectively. (B) Multiple sequence alignment of single- and double-RBEIII viral variants in respective subjects. A few TFBS of relevance are marked using open square boxes. The viral variants are aligned with the Indie.C1 reference sequence, which was pulsed in the sequencing sample as an internal control. Dashes represent sequence deletion and dots sequence homology. (C) Prognostic markers, plasma viral load (PVL) and CD4 cell count are represented by filled box with a solid line and open box with a dotted line, respectively.

## Discussion

The key finding of the present work is the continuing evolution of the HIV-1C viral promoter. In 2004, we reported the emergence of HIV-1C strains containing four copies of the NF-κB binding motif in the viral enhancer for the first time in India (14, 16). The 4-κB viral strains dominated the canonical viral strains containing three copies of NF-κB motifs in natural infection and under all experimental conditions, alluding to the additional copy of NF-κB motif conferring replication advantage (6). Subsequently, the 4-κB viral strains were also detected in Brazil and several African countries, suggesting that the phenomenon is not specific to a single country (15). Although our initial focus was limited to the NF-κB motif duplication and its impact on viral gene expression, we also observed the duplication of other TF binding motifs, including RBEIII and AP-1, though at a lower frequency (6, 16).

### Sequence motif duplication in HIV-1C differs from that of other HIV-1 families

Gene duplication accompanied by sequence variation played a crucial role in the acquisition of novel properties leading to the evolutionary success of organisms (20). In viruses, the duplication of biologically important sequence motifs may have provided the same survival advantage as gene duplication has done in higher organisms (18). The significance of sequence motif duplication in the coding sequences of HIV-1 has attracted more attention conventionally compared to that of regulatory sequences (21–23). Further, numerous publications have reported the deletion or duplication of different regions in the enhancer, core promoter, and modulatory regions that removed or added copies of TFBS (19, 24–26). However, such sequence modifications have been sporadic, typically limited to the individual or a small number of viral strains and cannot be generalized.

One notable exception to this observation is the occurrence of the MFNLP, which broadly represents the duplication of the RBEIII motif in the viral modulatory region with the concomitant co-duplication of the flanking sequences for other host factors. RBEIII motif duplication in HIV-1B was found in approximately 38% of primary viral isolates (8). Importantly, RBEIII motif duplication in HIV-1B is believed to ensure the presence of a binding site for RBF-2 when the original copy becomes non-functional due to mutations (27, 28).

RBEIII motif duplication in HIV-1C differs from that of HIV-1B in two crucial qualities. First, the creation of an additional RBEIII motif is not associated with the inactivation of the original motif. In other words, nearly all the double-RBEIII viral strains in our cohort of HIV-1C contained two copies of the intact motif without any mutations in the core sequence. Preliminary leads from our laboratory confirm the functional activity of both the motifs in such LTRs. Of note, the participants of the present study are all reportedly ART-naïve by self-declaration. We cannot rule out the possibility of ART exposure in HIV-1C, leading to the inactivation of the original RBEIII motif necessitating the need to create a second and functional RBEIII motif in the promoter. Second, the co-duplication of the RBEIII and NF-κB motifs is unique to HIV-1C, a property not seen in any other HIV-1 genetic subtype. Thus, HIV-1C appears to exploit the strategy of sequence motif duplication differently compared to other viral subtypes.

Importantly, the addition of more copies of NF-κB to the viral promoter may be beneficial by enhancing the transcriptional strength of the LTR. However, a stronger LTR can be detrimental to maintaining stable latency. HIV-1C appears to have found two different solutions to the paradox of gene expression regulation – limiting the copy number of the NF-κB motifs to three and duplicating the RBEIII motif.

### Limiting the number of NF-κB motif copy number in the viral enhancer

Three viral strains, LR-HHC, LR-FHHC, and LR-FHC, lack RBEIII duplication. The prevalence of the LR-FHHC viral strains was only 2% (13 of 607 primary viral isolates) in a southern Indian cohort when discovered during 2000-2003 for the first time (6). The prevalence of these strains increased to approximately 25% (39/159) during 2010-2011, evaluated at four different clinical sites of India, suggesting replication success of 4-κB viral strains at the population level (6). However, in the present study, the prevalence of the LR-FHHC viral strains dropped to 5.9% (27 of 455) during 2017-2019. Notably, a new variant viral strain LR-FHC representing the second-largest proportion among the emerging variants with 8.8% (40 of 455) was identified here for the first time. Given the reduction in the prevalence of LR-FHHC strains and the concomitant appearance of the LR-FHC strains, it is possible that the former is being replaced by the latter.

This observation leads to three logical conclusions. First, LR-FHHC strains, given the stronger transactivation properties of the promoter, may lack replication competence over a sustained period explaining the transient nature of their prevalence in the population. Second, the 4-κB viral strains must relinquish one κB-motif to regain the 3-κB formulation of the enhancer to down modulate the transcriptional strength of the viral promoter. The LR-FHHC strains relinquished one of the two H-κB sites to this end to become LR-FHC. Three, both the canonical LR-HHC and the variant LR-FHC strains contain the same number of NF-κB motifs in the enhancer. However, the three κB-motifs of the FHC-LTR are genetically variable. We propose that the LR-FHC-LTR is likely to be responsive to a broader range of cellular activation signals compared to the LR-HHC-LTR, given the NF-κB motif variation. Thus, by deleting one H-κB site from the LR-FHHC-LTR, HIV-1C appears to have down-modulated transcriptional strength of the viral promoter on the one hand but retained the broader reception potential to cellular signals on the other hand. If the LR-FHC viral strains enjoy a replication advantage at the population level, they are expected to replace the canonical HHC strains in the coming years.

### Is the RBEIII motif duplication to impose avid latency of a stronger viral promoter?

Seven different variant LTRs identified in this work contain a second copy of the RBEIII motif added by sequence-motif duplication (Fig.1 and 2*D-J*). Unlike in HIV-1B where a new RBEIII site is created as a compensatory mechanism when the original copy is mutated (8), in HIV-1C, both the RBEIII motifs are, in contrast, intact without a mutation in the core motif (5’-ACTGCTGA-3’). Thus, RBEIII duplication in HIV-1C appears to confer a novel function or an enhanced phenotype of the existing function but not compensating for a loss of function.

A second quality of the RBEIII motif duplication in HIV-1C is also relevant, especially for HIV latency. While RBEIII motif duplication is common to all the HIV-1 genetic subtypes (*SI Appendix*, Fig. S2 and see 7, 10, 11, 29), one significant distinction unique to HIV-1C is the co-duplication of the NF-κB motif, not seen in other subtypes. One variant LTR, LRhR-HHC, contains a total of four NF-κB motifs like the old FHHC-LTR. However, the variant LTR contains two RBEIII sites, unlike the FHHC-LTR that has only one (Fig. 1). Additionally, the duplicated κB-motif of LRhR-HHC is genetically distinct (5’-GGGACTTTC<u>A</u>-3’) from the other three types (C-, H-, and F-κB) described above. The Single Nucleotide Mutation Model predicted the 5’-GGGACTTTC<u>A</u>-3’ motif to bind the p50 homodimer with reduced affinity compared to the consensus NF-κB motif 5’-GGGACTTTCC-3’. This binding prediction was supported by the bimolecular dsDNA microarray analysis (30). Lastly, the duplicated κB-motif of LRhR-HHC-LTR is separated from the viral enhancer by one RBEIII motif thus, obliterating the distinction between the viral modulatory and enhancer elements. The biological significance of creating one of each RBEIII and NF-κB motifs in the LRhR-HHC-LTR is of interest.

In the absence of cell activation, the RBEIII motif functions predominantly as a repressive element by recruiting RBF-2 comprising three different cell factors, including TFII-I (27). While TFII-I can activate several cellular genes, it can also suppress gene expression from several other cellular promoters, including c-*fos* (31). Thus, the presence of two copies of the RBEIII motif in the LTR may have a profound impact on viral latency, probably by stabilizing the latency phase. The NF-κB binding motifs, in contrast, play a predominantly positive role in enhancing transcription from the LTR, under the conditions of cell activation. Thus, a higher copy number of NF-κB (4 copies Vs. 3) in the promoter may offset the negative impact of the RBEIII motifs, especially when the provirus is induced out of latency. Of note, unlike the variant LRhR-HHC, a different variant strain LRhR-HC contains one less NF-κB motif (two RBEIII but only three NF-κB motifs). Preliminary results from our laboratory show that the LRhR-HC-LTR requires a profoundly stronger activation signal, compared to LRhR-HHC-LTR or the canonical LR-HHC-LTR, for latency reversal in Jurkat cells or primary CD4 cells (Bhange D et al, unpublished observations).

A different variant promoter pair (LRLR-HHC and LRLR-HC) is also of interest in this respect. This pair also contains an RBEIII motif duplication where the motifs are accompanied by the co-duplication of the TCF-1α/LEF-1 motif, not the NF-κB site. One member of the pair contains three NF-κB motifs (LRLR-HHC) while the other only two (LRLR-HC). This variant promoter pair may have similar gene expression properties as that of the LRhR-HHC and LRhR-HC LTRs if the additional copy of TCF-1α/LEF-1 is a functional equivalent of the NF-κB motif.

### The implication of promoter variation on HIV-1 pathogenesis and evolution

Since a single promoter regulates the expression of all the HIV-1 proteins and controls latency, a profound variation in the TFBS composition is expected to have a significant impact on the various properties of the virus, including latency, viral load, disease progression, and viral evolution. The evolution of regulatory elements may play a role as essential or even more important than that of coding sequences (20). However, little attention was focused on the evolution of the regulatory elements in HIV-1, unlike the protein-coding regions (32).

In our study, the cross-sectional and longitudinal analyses did not find a statistically significant difference in the levels of any of the prognostic markers across the promoter variants categorized into four groups, HHC, FHC, FHHC, and RR (Fig. 4). A significant difference in PVL and CD4 cell count was found in a previous study from our laboratory when a cohort of eighty patients was divided into HHC and FHHC groups (6). The present study did not find such differences, except for the RR group manifesting a trend in the complete-case analysis, which was not statistically significant. The analytical power of the present study was profoundly compromised given the loss of available samples due to the implementation of the test-and-treat policy.

Commensurate with our findings, earlier studies of the RBEIII motif duplication in HIV-1B also failed to see an association between the LTR profile and clinical or transcriptional phenotypes in a cross-sectional cohort (8). A different study could not see a correlation between the RBEIII motif duplication and syncytium-inducing property of envelope and, thus, disease progression (33). Likewise, Koken S., et al. demonstrated the domination of a viral strain containing a single copy of the RBEIII motif over a counterpart containing two copies of the TFBS in 28 days in cell culture. However, no such differences were observed in patients alluding to an association between the RBEIII copy-number and disease progression (33).

The coexistence of viral strains could be a likely explanation for the absence of association between LTR variant forms and prognostic markers in our cohort. Deep sequencing of the samples identified the presence of a co-infection in all four subjects in our study. At the current time, it appears that the double-RBEIII viral strains appear only as a co-infection along with the single-RBEIII strains. In three of the four subjects, single-RBEIII viral strains seem to dominate the double-RBEIII variants in both the genomic DNA and RNA compartments and at most of the follow-up time-points. If RBEIII duplication indeed manifests a suppressive effect on viral gene expression, a distinct association between the duplication and the prognostic markers may become evident in a mono-infection, but not in a coinfection. The dominant influence of the RBEIII motif duplication on viral gene expression has been confirmed using panels of engineered viral clones (Bhange D et al, unpublished observations).

In summary, our work records the emergence of several promoter variant viral strains in HIV-1C of India over recent years. Sequences representing the variant viral forms are also found in the sequence databases derived from different global regions where HIV-1C is predominant. Sequence motif duplication creates additional copies of TFBS that play a crucial role in regulating HIV latency and even blurs the distinction between the viral enhancer and modulatory regions. Given that the RBEIII and AP-1 sites play a significant role in regulating latency (19 and 20), the influence of RBEIII site duplication, especially when accompanied by the co-duplication of NF-κB motifs, needs experimental evaluation. Consistent monitoring will be necessary to understand which variant viral strains will survive to establish spreading epidemics in coming years. Detailed investigations are warranted to evaluate the impact of the TFBS profile differences on HIV-1 latency and latent reservoir properties. ART administration may have a profound impact on the promoter variations described here by exacerbating such sequence duplications.

## Materials and methods

### Study participants and samples

Participants were recruited at four different sites in India for primary screening (PS) and longitudinal study (LS)-All India Institute of Medical Sciences (AIIMS), New Delhi (PS=107, LS=73); National AIDS Research Institute (NARI), Pune (PS=61, LS=38); St. John’s Medical Hospital, Bangalore (PS=116, LS=60); and Y. R. Gaitonde Centre for AIDS Research and Education (YRG CARE), Chennai (PS=171, LS=37). Subjects above 18 years of age with a documented evidence of serological positive test for HIV-1 were recruited to the study. All the study subjects were ART naïve at baseline as per self-reporting.

### Ethics statement

Written informed consent was obtained from all study participants, following specific institutional review board-approved protocols. Ethical approval for the study was granted by the Institutional Review Board of each clinical site. All the clinical sites screened the potential subjects, counseled, recruited the study participants, and maintained the clinical cohorts for the present study. The Human Ethics and Biosafety Committee of Jawaharlal Nehru Centre for Advanced Scientific Research (JNCASR), Bangalore, reviewed the proposal and approved the study.

### Primary screening: LTR amplification and molecular typing of the viral promoter

For the molecular typing of the viral promoter, 15 ml of peripheral blood were collected from every participant at one time. Genomic DNA was extracted from 200 µl of the whole blood, and the U3 region of LTR was amplified using a nested-PCR strategy (details in *SI Appendix*). The amplified LTR sequences were analyzed using Sanger Dideoxy sequencing and were subjected to further quality control by multiple sequence alignment and phylogenetic analysis with the in-house laboratory sequence database using the ClustalW algorithm of BioEdit sequence alignment editor and MEGA6.0 software, respectively. Different viral strains were categorized by analyzing the LTR spanning modulatory and enhancer region from the LEF motif up to the Sp1III motif.

### Phylogenetic analysis

The phylogenetic analysis of the HIV-1C LTR variants derived from 461 patient samples was performed using MEGA7.0 software. The analysis was performed with 1,000 bootstrap values. The percentage of clustering is shown using the Maximum Composite Likelihood method. The values presented represent the number of base substitutions per site. All the positions containing gaps and missing data were eliminated. A total of 412 positions were included in the final dataset.

### The follow-up clinical procedures

Following successful characterization of the viral promoter at JNCASR, the clinical sites were advised to recruit specific study subjects without disclosing the nature of the viral LTR. The clinical sites were, thus, blinded to the identity of the viral LTR. All the clinical procedures were performed at the clinical sites using the same protocols and kits as described in *SI Appendix*. The analysis of CD4 cell count, plasma viral load, and soluble CD14 was performed at longitudinal time-points (details in *SI Appendix*).

### RNA isolation and RT-PCR for the next-generation sequencing

RNA was extracted from 1 ml of the stored plasma samples, and the complementary DNA (cDNA) was synthesized using HIV-specific primers listed in the *SI Appendix* Table S1 (details in *SI Appendix*). The cDNA was used for the amplification of LTR.

### The next-generation sequencing

The PCR products containing the RBEIII motif duplication were subjected to the NGS analysis using the Miseq Illumina platform. Each sample was amplified in duplicates using primers containing a unique 8 bp barcode sequence specific for each sample. The amplification of the U3 region (∼300-350 bp) using genomic DNA or cDNA prepared from plasma RNA was performed the same way as described for the primary screening except that the primers contained a unique sequence barcode at the 5’-end as listed in *SI Appendix*, Table S1 and 2. The concentration of the purified PCR product was determined using the Qubit™ dsDNA BR assay kit, and all the samples were pooled at an equal concentration and were processed further. The final sample included a pulsed LTR amplicon of Indie-C1, a reference HIV-1C molecular clone, as internal quality control for sequencing. The pooled sample was further processed for Illumina MiSeq sequencing (details in *SI Appendix*).

Data analysis was performed using a pipeline as depicted (*SI Appendix*, Fig. S4) and the details are mentioned in *SI Appendix*. Following the analysis procedure, the percentage prevalence of single and double RBEIII variants was obtained (Fig. 5).

### Statistical analysis

The data were analyzed using GraphPad prism 9, except the sequences used for phylogeny determination. P values of 0.05 or less were considered statistically significant. A nonparametric, Kruskal–Wallis (for multiple comparisons) test was applied to evaluate statistical significance in the case of a cross-sectional analysis of plasma viral load (Fig. 4*A* and *SI Appendix*, Fig. S3*A*). One-way ANOVA was used to evaluate statistical significance for cross-sectional analysis of CD4 cell count and sCD14 levels (Fig. 4*B, C* and *SI Appendix*, Fig. S3*B, C)*. Two-way ANOVA was used to evaluate statistical significance in the case of longitudinal analysis of all the three parameters, plasma viral load, CD4 cell count, and sCD14. (Fig. 4, and *SI Appendix*, Fig. S3).

## Data availability statement

All the sequences reported in this paper are available from the GenBank database under accession nos. MN840242.1-MN840356.1, MT847032 -MT847207, MT593868 -MT594037. The raw data files for Illumina MiSeq are available under accession no. PRJNA720640.

## Acknowledgments and funding sources

We thank Dr. Kushgra Bansal for the discussions during the NGS data analysis. We thank the study participants for their participation and cooperation during the complete duration of the clinical study. This work was supported by the Department of Biotechnology, Ministry of Science and Technology, Government of India (Sanction order no. BT/PR7359/MED/29/651/2012).

## Abbreviations

TFBS: Transcription factor binding site,
HIV-1C: HIV-1 subtype C,
LTR: Long terminal repeats,
C-LTR: HIV-1C LTR,
NGS: Next-generation sequencing,
RT-PCR: Reverse transcription polymerase chain reaction,
ART: Antiretroviral therapy.

