## Supplementary information for "The evolution of regulatory elements in the emerging promoter variant strains of HIV-1"

### SI Appendix

**This PDF file includes:**

Extended methods

Figures S1 to S4

Tables S1 and S3

#### Methods and materials

##### Primary screening: LTR amplification and molecular typing of the viral promoter

To type the viral promoter, 15 ml of peripheral blood was collected from every participant at one time. Of this sample, 3-5 ml of blood was allocated to determine the CD4 cell count by flow cytometry and the extraction of genomic DNA from whole blood. From the rest of the blood sample, PBMC were isolated by density-gradient centrifugation. The PBMC and plasma samples were stored in 1 ml aliquots in a liquid nitrogen container or a deep freezer, respectively, for further clinical analysis.

Genomic DNA was extracted from 200 µl of the whole blood using a commercial DNA extraction kit (GenElute™ Blood Genomic DNA kit, Cat. No. NA2020, Sigma Aldrich, U.S.A.) and was eluted in 200 µl volume. The extracted DNA samples from the clinical sites were shipped to JNCASR on cold packages. The LTR sequences were amplified using 200-300 ng of genomic DNA as a template from each of the clinical samples using Taq DNA Polymerase (Cat. No. M073L, New England Biolabs) in a Peqstar 2x thermal cycler (Peqlab, VWR). The U3 region of LTR was amplified using a nested-PCR strategy with the primers listed in the *SI Appendix,* Table S1. Strict procedural and physical safeguards were implemented to minimize carryover contamination that included reagent preparation and PCR setup, amplification, and post-PCR processing of samples in separate rooms. The PCR products were purified using a commercial DNA purification kit (FavorPrep Gel/PCR Purification Kit, Cat. No. FAGCK001, Favorgen Biotech Corp. Ping-Tung 908, Taiwan). The amplified LTR sequences were analyzed using Sanger Dideoxy sequencing (Applied Biosystems, California, U.S.A.). All the sequences were subjected to further quality control by multiple sequence alignment and phylogenetic analysis with the in-house laboratory sequence database using the ClustalW algorithm of BioEdit sequence alignment editor and MEGA6.0 software, respectively. The modulatory-and-enhancer region of the LTR spanning from the LEF motif up to the Sp-1III motif was analyzed to type the viral strains.

##### The follow-up clinical procedures

Following successful characterization of the viral promoter at JNCASR, the clinical sites were advised to recruit specific study subjects without disclosing the nature of the viral LTR. The clinical sites were, thus, blinded to the identity of the viral LTR. All the clinical procedures were performed at the clinical sites using the same protocols and kits as described below.

A total of 15 ml of peripheral blood was collected in a BD vacutainer (Cat. no. 367525, Becton Dickinson, California, U.S.A.) from each participant at 0, 6, 12, 18, 24, and 36-months during 2015-19. The PBMC and plasma samples were stored in 1 ml aliquots in a liquid nitrogen container or a deep freezer, respectively. The CD4 T-cell count was determined using the BD Multitest commercial kit-CD3/CD8/CD45/CD4 (Cat. No. 340491, Becton Dickinson, California, U.S.A.) following the manufacturer's instructions. The samples were analyzed using a BD FACS Calibur flow cytometer or any other suitable machine. Calibration of the flow cytometer was performed using BD CaliBRITE 3 and APC beads (Cat. no. 340486 and 340487, respectively, Becton Dickinson, California, U.S.A.). The plasma viral RNA load was determined at 0 and 12-month time-points using the Abbott m2000rt Real-Time PCR machine (Abbott Molecular Inc. Des Plaines, IL, U.S.A.). Levels of soluble CD14 (sCD14) in the plasma were determined using Human sCD14 Quantikine ELISA Kit (Cat No. DC140, R & D Systems, Minnesota, U.S.A.). Analysis of sCD14 was performed at months 0 and 12.

##### RNA isolation and RT-PCR

RNA was extracted from 1 ml of the stored plasma samples using a commercial Viral RNA isolation kit (the NucliSENS miniMAG nucleic acid extraction kit, Ref. No. 200293, BioMerieux, France). The complementary DNA (cDNA) was synthesized using HIV-specific primers (Table S1) and a commercial kit (SuperScript™ IV First-Strand Synthesis System with ezDNase™ Enzyme (Cat. No. 18091150), Invitrogen, Carlsbad, California, U.S.A.). The reaction vials were incubated at 65°C for 5 min, following 2 min incubation on ice and 50°C for 50 min. The reactions were terminated by incubating the samples at 85°C for 5 min followed by RNaseH treatment. The cDNA was used for the amplification of LTR.

**The next-generation sequencing**

The PCR products containing the RBEIII motif duplication were subjected to the NGS analysis using the Miseq Illumina platform. Each sample was amplified in duplicates using primers containing a unique 8 bp barcode sequence specific for each sample. The amplification of the U3 region (~300-350 bp) using genomic DNA or cDNA prepared from plasma RNA was performed the same way as described above for the primary screening except that the primers contained a unique sequence barcode at the 5'-end as listed (*SI Appendix,* Tables S1, and S2). The concentration of the purified PCR product was determined using the Qubit™ dsDNA BR assay kit (Cat. No. Q32850, Invitrogen, California, United States). All the samples were pooled at an equal concentration and were processed further. We pulsed the LTR amplicon of Indie-C1, a reference HIV-1C molecular clone, as internal quality control for sequencing.

DNA was quantified using the QUBIT 3 Fluorometer and a dsDNA HS Dye. After adding an 'A' nucleotide to the 3' ends, the adenylated fragments were ligated with loop adapters and cleaved with uracil-specific excision reagent (USER) enzyme. The DNA was further purified using AMPure beads and then enriched by PCR in 6 cycles using NEBNext Ultra II Q5 master mix (Cat. No. E7645L, New England Biolabs, Inc., Massachusetts, U.S.A.), Illumina universal primer, and sample-specific octamer primers. The amplified products were cleaned by using AMPure beads, and the final DNA library was eluted in 15 µl of 0.1X TE buffer. The volume of 1 µL of the library was used to quantify by QUBIT 3 Fluorometer using dS DNA HS reagent. The fragment analysis was performed on Agilent 4150 Tape Station by loading 1 µl of the library to Agilent D1000 Screen Tape. The library was sequenced using the Illumina MiSeq system and MiSeq Reagent kit v3 (Cat. No. MS-102-3003, Illumina, San Diego, CA, U.S.A.) following the manufacturer's instructions.

Data analysis was performed using a custom pipeline as depicted (*SI Appendix,* Fig. S4). First, the quality assessment was performed using FastQC (version 0.11.5), and the sequencing adapters were removed using Trimmomatic-master (version 0.33) from the raw paired-end data. Second, the paired reads from both the forward and reverse files were merged using the PEAR algorithm of PANDAseq with the minimum and maximum read length set to be 200 and 500, respectively, with an overlap of at least 8 base pairs. Next, the merged reads were mapped to the LTR region of the two reference sequences: Indie.C1 (AB023804) and D24 (EF469243.2), using local alignment with Bowtie2 software. Using custom c++ and shell scripts, the mapped reads were demultiplexed in individual samples based on the combination of forward and reverse barcodes (*SI Appendix,*Table S2). The reads containing the C-κB motif sequence (HIV-1C) were considered for further analysis using a custom c++ script. All the HIV-1C reads were then grouped in multiple categories based on the number and sequence of NF-κB, RBEIII, and TCF-1α /LEF-1 motifs using a custom c++ script. The percentage of every category was then calculated based on the total number of reads within each sample, using the custom shell, c++, and R scripts. Next, DeconSeq (version 0.4.3) was used to analyze and filter out inter-sample sporadic contamination. A reference database was prepared manually for every major-variant category (category with >10% of total reads at all time points), by taking only the most abundant variant of that category from each time-point of the sample. The reference database of a variant category was used to cross-check for contamination in the same variant category of the other samples, where the variant category is <10% in one or more time-points. The percent coverage and identity thresholds were set to 90% each to allow a maximum of 5 mismatches or indels in a sequence stretch of 50 bases. The sum of percentages of all the variants classified as clean for a category, in all the time-points, was done using custom shell and R scripts. From the total number of reads for each time-point of a sample, the number of contaminated reads was eliminated, and the clean reads were taken ahead for calculating the percentage prevalence of single and double RBEIII variants.

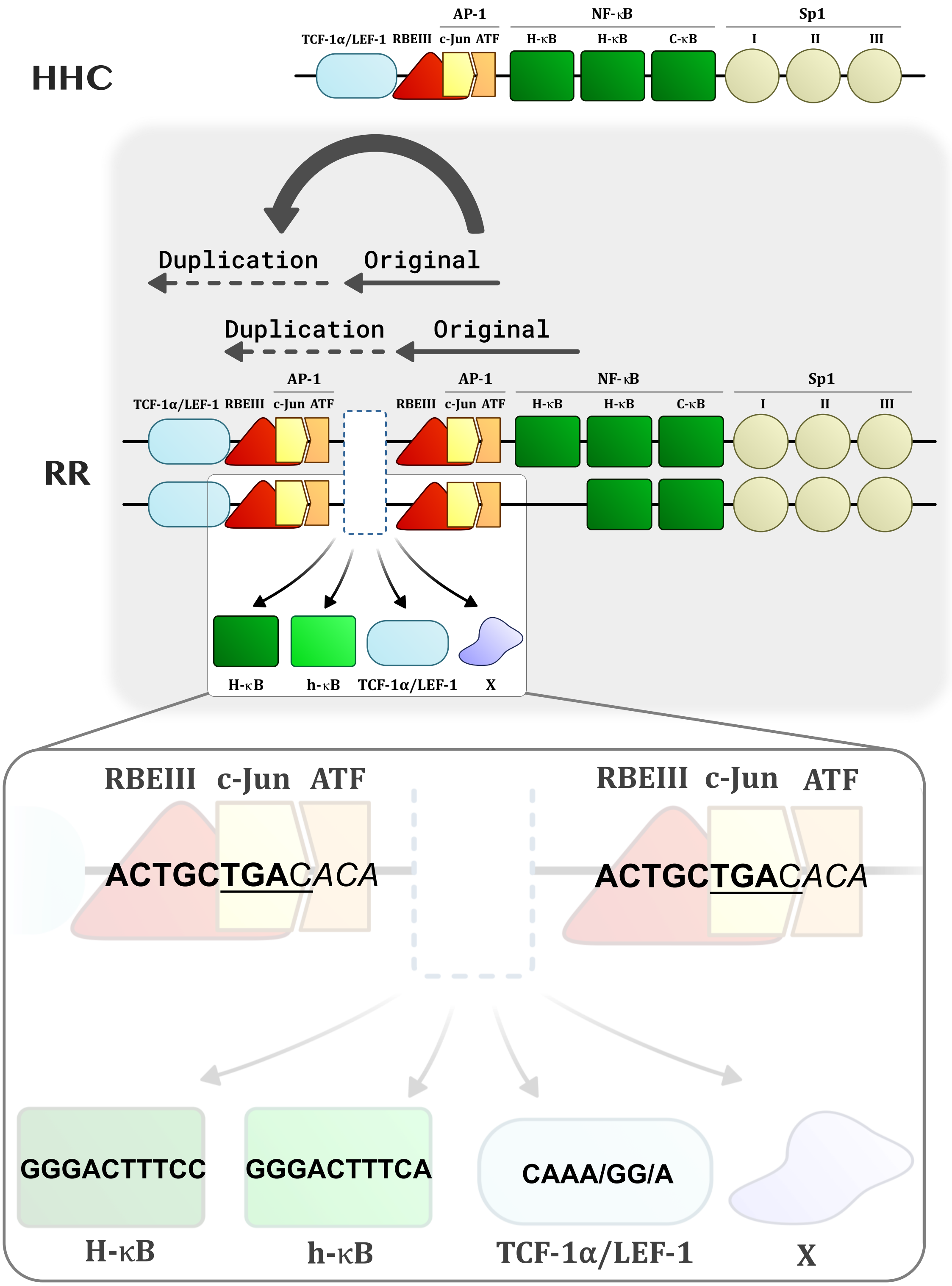


**Fig. S1.** A schematic of duplication pattern in RR group of emerging HIV-1C variants. A top panel shows the arrangement of TFBS in the HHC variant. Middle panel depicts the variation in the number and position of TFBS in the RR group of variants. The RBF-2, c-Jun and ATF (together termed as AP-1) binding motifs appear as a cluster which is shown to be duplicated in the RR variants. The original and duplicated region of TFBS is marked with solid and dotted line, respectively. The region between two RBEIII-AP-1 complex is depicted with white dotted-arrow which suggest the variation of the TFBS which are further illustrated at the bottom of the second panel. The last panel focuses on the nucleotide sequence of the RBEIII-AP-1 complex along with the sequence intervening the two RBEIII-AP-1 clusters.

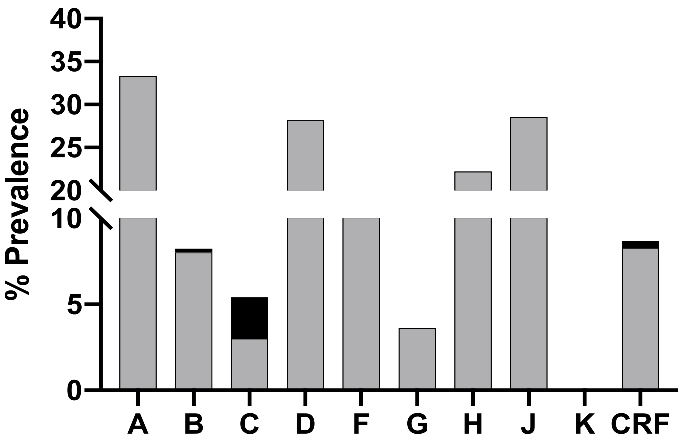


| | A | B | C | D | F | G | H | J | K | CRF |
|---|---|---|---|---|---|---|---|---|---|---|
| n | 9 | 1395 | 831 | 85 | 74 | 83 | 9 | 7 | 3 | 2168 |
| RBEIII & NF-κB | 0 | 3 | 20 | 0 | 0 | 0 | 0 | 0 | 0 | 8 |
| Only RBEIII | 3 | 112 | 25 | 24 | 9 | 3 | 2 | 2 | 0 | 180 |

**Fig. S2.** The relative prevalence of the RBEIII and NF-κB motif duplication in HIV-1 subtypes. All the available LTR sequences containing the duplication of one or both TFBS were downloaded from the LANL HIV database and categorized under each major subtype as shown (n). A single sequence per patient was included in the analysis. The grey bar represents the prevalence of viral sequences containing the duplication of RBEIII motif and the filled bar that of both RBEIII and NF-κB motifs.

**A** Plasma viral load

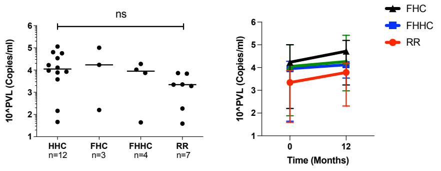


**Sample size and statistics**

| Variant | M0 | | | M12 | | |
|---|---|---|---|---|---|---|
| | n | Median | Percentile (10 - 90%) | n | Median | Percentile (10 - 90%) |
| HHC | 12 | 11,203.0 | 76.0 - 1,00,056.0 | 12 | 18,450.0 | 952.3 - 2,62,976.0 |
| FHC | 3 | 17,157.0 | 159.0 - 1,01,041.0 | 3 | 52,539.0 | 1,725.0 - 1,56,000.0 |
| FHHC | 4 | 8,964.0 | 44.0 - 18,868.0 | 4 | 13,266.0 | 3,468.0 - 18,240.0 |
| RR | 7 | 2,216.0 | 39.0 - 7,287.0 | 7 | 6,090.0 | 204.0 - 15,800.0 |

**B** CD4 cell count

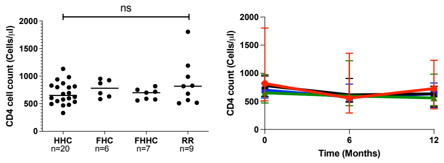


| Variant | M0 | | | M6 | | | M12 | | |
|---|---|---|---|---|---|---|---|---|---|
| | n | Median | Percentile (10 - 90%) | n | Median | Percentile (10 - 90%) | n | Median | Percentile (10 - 90%) |
| HHC | 20 | 650.5 | 470.2 - 977.9 | 20 | 595.5 | 422.1 - 1223.0 | 20 | 561.5 | 423.5 - 983.6 |
| FHC | 6 | 780.0 | 584.0 - 953.0 | 6 | 616.0 | 485.0 - 908.0 | 6 | 634.0 | 402.0 - 769.0 |
| FHHC | 7 | 700.0 | 559.0 - 818.0 | 7 | 577.0 | 499.0 - 805.0 | 7 | 632.0 | 389.0 - 827.0 |
| RR | 9 | 817.0 | 509.0 - 1804.0 | 9 | 557.0 | 292.0 - 1356.0 | 9 | 727.0 | 367.0 - 1,231.0 |

**C** sCD14

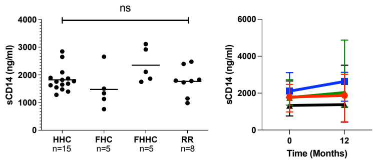


| Variant | M0 | | | M12 | | |
|---|---|---|---|---|---|---|
| | n | Media n | Percentile (10 - 90%) | n | Median | Percentile (10 - 90%) |
| HHC | 15 | 1,755.0 | 1,349.0 – 2,727.0 | 15 | 2,033.0 | 1,252.0 – 4,872.0 |
| FHC | 5 | 1,333.0 | 763.0 – 2,655.0 | 5 | 1,376.0 | 428.8 – 3,514.0 |
| FHHC | 5 | 2,108.0 | 1,748.0 – 3,110.0 | 5 | 2,631.0 | 1,560.0 – 3,128.0 |
| RR | 8 | 1,785.0 | 983.0 – 2,474.0 | 8 | 1,874.0 | 448.3 – 3,015.0 |

**Fig. S3.** Cross-sectional and longitudinal analyses of prognostic markers under the 'complete-case' scenario. Plasma viral load (A), CD4 cell count (B), and soluble CD14 (C) levels of the four study arms are presented at the baseline (left panels) and follow-up points (right panels). The number of samples included under each evaluation and the corresponding statistics are presented in the tables. Given the limited sample numbers, several groups were pooled under the double-RBEIII arm. A non-parametric test i.e. Kruskal-Wallis test was applied for the statistical analysis of the plasma viral load. One-way ANOVA with Dunnet multiple comparison test was applied to CD4 count and sCD14 . Two-way ANOVA was used for the comparison of the longitudinal analysis.

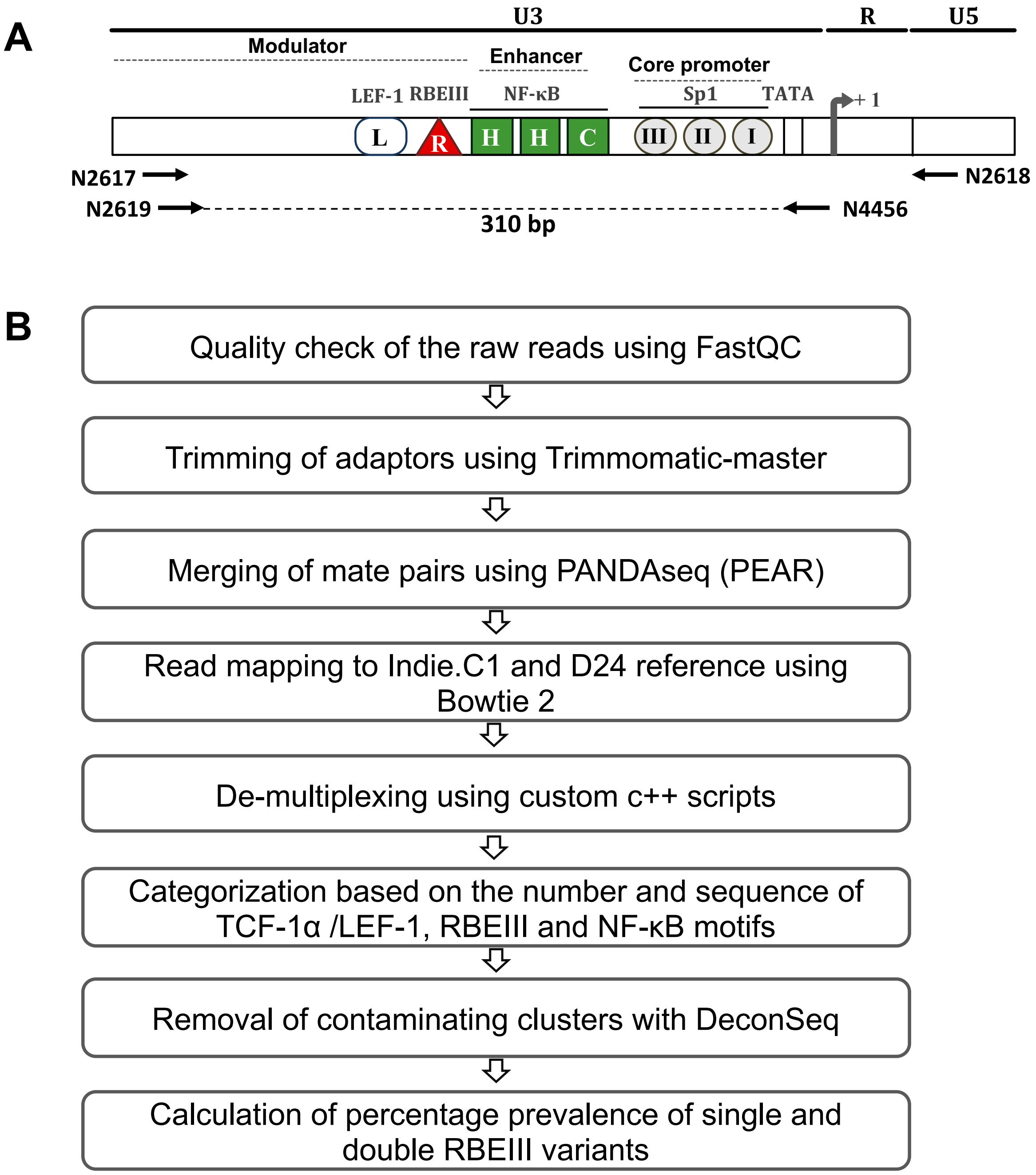


**Fig. S4.** A schematic representation of the strategy for PCR amplification and Illumina sequencing of the LTR. (A) The location and orientation of the primers amplifying the LTR region are depicted. A PCR fragment of 310 bp was used for the NGS. (B) The PCR fragments were analysed using the pair-end MiSeq Illumina sequencing. The flow-chart depicts the pipeline followed for data analysis. The reads were mapped to the reference sequence of pIndie.C1.

**Table S1**: Primers used for PCR and cDNA synthesis

| Purpose | | Primer no. | Sequence (5′-3′) |
|---|---|---|---|
| **Primary screening** | OFP | N2617M | CCACTGACCTTTGGATGGTGCTTCAAG |
| | ORP | N2618 | TGAGGGATCTCTAGTTACCAGAGTCAC |
| | IFP | N2619 | TGCTTCAAGCTAGTRCCAGTTGACCCA |
| | IRP | N2620 | AGAGTCACACAACAGACGGGCACACA |
| **Sanger sequencing** | | N2621 | AAGCTGGTACCAGTTGAC |
| **cDNA synthesis** | | N4101 | TTTTTTTTTTTTTTCAGAGCACTC |
| | | N4102 | TTTTTTTTTTTTTCAGAGCACTCAAG |
| | | N4103 | TTTTTTTTTTTTTTCAGAGCACTCAAGG |
| **NGS** | OFP | N2619 | TGCTTCAAGCTAGTRCCAGTTGACCCA |
| | ORP | N2618 | TGAGGGATCTCTAGTTACCAGAGTCAC |
| | IFP | N2619BX | (8 bp barcode)-TGCTTCAAGCTAGTRCCAGTTGACCCA |
| | IRP | N4456BX | (8 bp barcode)- GCGAAAAGCAGCTGCTTATATGCAGCATCTGAGGG |

OFP= Outer forward primer, ORP= Outer reverse primer, IFP= Inner forward primer, IRP= Inner reverse primer, X= Number of the barcode

**Table S2**: (A) Combination of barcoded primers used for LTR amplification from each sample, (B) Sequence of each 8 bp barcode

**A**

| Sample No. | Month | Genomic DNA | | Plasma RNA | |
|---|---|---|---|---|---|
| | | Set 1 | Set 2 | Set 1 | Set 2 |
| **2079** | M0 | B5 F | B6 F | B5 F | B6 F |
| | | B1 R | B1 R | B12 R | B12 R |
| | M6 | B5 F | B6 F | B5 F | B6 F |
| | | B4 R | B4 R | B13 R | B13 R |
| | M12 | B5 F | B6 F | B5 F | B6 F |
| | | B7 R | B7 R | B15 R | B15 R |
| **4084** | M0 | B1 F | B2 F | B1 F | B2 F |
| | | B4 R | B4 R | B7 R | B7 R |
| | M6 | B1 F | B2 F | B1 F | B2 F |
| | | B5 R | B5 R | B11 R | B11 R |
| | M12 | B1 F | B2 F | B1 F | B2 F |
| | | B6 R | B6 R | B12 R | B12 R |
| **3767** | M0 | B3 F | B4 F | B3 F | B4 F |
| | | B5 R | B5 R | B11 R | B11 R |
| | M6 | B3 F | B4 F | B3 F | B4 F |
| | | B6 R | B6 R | B12 R | B12 R |
| | M12 | B3 F | B4 F | B3 F | B4 F |
| | | B7 R | B7 R | B13 R | B13 R |
| **VFSJ20** | M0 | B13 F | B14 F | B13 F | B14 F |
| | | B19 R | B19 R | B1 R | B1 R |
| | M12 | B13 F | B14 F | B13 F | B14 F |
| | | B20 R | B20 R | B4 R | B4 R |
| | M24 | B13 F | B14 F | B13 F | B14 F |
| | | B22 R | B22 R | B7 R | B7 R |

**B**

| Barcode No. | Sequence (5'-3') |
|---|---|
| B1 | TCGCCTTA |
| B2 | CTAGTACG |
| B3 | TTCTGCCT |
| B4 | GCTCAGGA |
| B5 | AGGAGTCC |
| B6 | CATGCCTA |
| B7 | GTAGAGAG |
| B9 | TGCCTCTT |
| B10 | TCCTCTAC |
| B11 | TCATGAGC |
| B12 | CCTGAGAT |
| B13 | TAGCGAGT |
| B14 | GTAGCTCC |
| B15 | TACTACGC |
| B19 | GAGCGCTA |
| B20 | CGCTCAGT |
| B22 | ACTGATCG |

**Table S3**: Proportion of promoter variants at each clinical site

| Category | Variant | AIIMS | NARI | St. John's Hospital | YRG CARE | All clinics |
|---|---|---|---|---|---|---|
| I | HHC | 67 (62.6%) | 38 (62.3%) | 79 (68.1%) | 119 (69.6%) | 303 (66.6%) |
| | FHC | 13 (12.2%) | 5 (8.2%) | 8 (6.9%) | 14 (8.2%) | 40 (8.8%) |
| II | FHHC | 7 (6.5%) | 6 (9.8%) | 9 (7.8%) | 5 (2.9%) | 27 (5.9%) |
| III | RR | 20 (18.7%) | 12 (19.7%) | 20 (17.2%) | 33 (19.3%) | 85 (18.7%) |
| **Total** | | **107** | **61** | **116** | **171** | **455** |
